# Phosphorylation of VapB antitoxins affects intermolecular interactions to regulate VapC toxin activity in *Mycobacterium tuberculosis*

**DOI:** 10.1101/2024.05.30.596101

**Authors:** Basanti Malakar, Valdir Barth, Julia Puffal, Nancy Woychik, Robert N. Husson

**Author notes:** Address correspondence to Basanti Malakar, or Robert N. Husson. Valdir Barth, Institute for Biomedical Sciences, University of São Paulo, São Paulo, Brazil. Julia Puffal, USACE Engineer Research and Development Center, Biotechnology Research Facility, Austin, TX, Bennett Aerospace Inc, Raleigh, NC, University of Texas, Defense Research Advancement, Austin, TX.

## Abstract

Toxin-antitoxin modules are present in many bacterial pathogens. The VapBC family is particularly abundant in members of the *Mycobacterium tuberculosis* complex, with 50 modules present in the *M. tuberculosis* genome. In type IIA modules the VapB antitoxin protein binds to and inhibits the activity of the co-expressed cognate VapC toxin protein. VapB proteins also bind to promoter region sequences and repress expression of the *vapB-vapC* operon. Though VapB-VapC interactions can control the amount of free VapC toxin in the bacterial cell, the mechanisms that affect this interaction are poorly understood. Based on our recent finding of Ser/Thr phosphorylation of VapB proteins in *M. tuberculosis*, we substituted phosphomimetic or phosphoablative amino acids at the phosphorylation sites of two VapB proteins. We found that phosphomimetic substitution of VapB27 and VapB46 resulted in decreased interaction with their respective cognate VapC proteins, whereas phosphoablative substitution did not alter binding. Similarly, we determined that phosphomimetic substitution interfered with VapB binding to promoter region DNA sequences. Both decreased VapB-VapC interaction and decreased VapB repression of *vapB*-*vapC* operon transcription would result in increased free VapC in the *M. tuberculosis* cell. *M. tuberculosis* strains expressing *vapB46-vapC46* constructs containing a phosphoablative *vapB* mutation resulted in lower toxicity compared to a strain expressing native *vapB46*, whereas similar or greater toxicity was observed in the strain expressing the phosphomimetic *vapB* mutation. These results identify a novel mechanism by which VapC toxicity activity can be regulated by VapB phosphorylation, potentially in response to extracytoplasmic as well as intracellular signals.

**Importance:** Intracellular bacterial toxins are present in many bacterial pathogens and have been linked to bacterial survival in response to stresses encountered during infection. The activity of many toxins is regulated by a co-expressed antitoxin protein that binds to and sequesters the toxin protein. The mechanisms by which an antitoxin may respond to stresses to alter toxin activity are poorly understood. Here we show that antitoxin interactions with its cognate toxin, and with promoter DNA required for antitoxin and toxin expression, can be altered by Ser/Thr phosphorylation of the antitoxin, and thus affect toxin activity. This reversible modification may play an important role in regulating toxin activity within the bacterial cell in response to signals generated during infection.

## Introduction

There were an estimated 10.6 million new cases of tuberculosis (TB) disease in 2022 and approximately a quarter of the world’s population is thought to have been infected with *Mycobacterium tuberculosis* (1). Though a small proportion of those who are infected develop TB disease weeks to months after infection, most people control the infection, and many may eradicate it while retaining persistent immunoreactivity to *M. tuberculosis* antigens (2). In a substantial minority of cases, however, the infection may be controlled without being eliminated, resulting in asymptomatic (latent) TB infection. The development of TB disease and the processes that lead to bacterial eradication versus persistence following initial infection, as well as reactivation to cause late onset TB, all require regulation of bacterial growth and metabolism in response to host factors. During TB disease, for example, there are periods of rapid replication, e.g. following initial infection, and restricted replication, e.g. in mature granulomas or in centers of caseous necrosis.

Though bacterial replication is essential for infection and dissemination within the host, within populations of growing *M. tuberculosis* and other bacteria, there is a subset of cells that are not actively growing (3–6). These persister cells are phenotypically tolerant to antibiotics and to host bactericidal stresses and can resume growth when the stress is no longer present.

Imaging data have shown striking heterogeneity of TB lesions in individual animals and humans with TB disease or early latent TB, and of *M. tuberculosis* cells, with evidence of hypoxia and dormancy regulon activation in bacteria within TB lesions (7–10). These features of TB pathogenesis highlight the importance of understanding *M. tuberculosis* metabolic and growth adaptations and the ways they are regulated in response to environmental stimuli.

Toxin-Antitoxin (TA) systems are present in a broad range of bacteria, including many human pathogens (11, 12). In contrast to secreted bacterial toxins that disrupt host functions, the toxin proteins of TA systems typically remain intracellular, where they modulate bacterial physiology. Although other activities have been identified, most TA toxins have been found to cleave RNAs, including mRNAs, tRNAs and rRNAs, resulting in translation inhibition, proteome remodeling, and reversible growth arrest (12–18). TA toxins have been shown to enhance bacterial survival in the setting of several stress conditions (14).

*M. tuberculosis* has more than 80 TA systems, far more than most bacteria including other mycobacteria, suggesting a role for these systems in fine tuning growth and *M. tuberculosis* physiology in response to the variety of conditions that are encountered during infection (18). These proteins have been hypothesized to play a role in TB pathogenesis, including antimicrobial tolerance and long-term persistence within granulomas during latent TB infection (18–20). Some toxins may also engage in the regulation of host-generated stresses such as oxidative or nitrosative stress and consequently play a role in controlling bacterial pathogenesis (21).

Type II TA systems, which comprise most of the TA systems present in *M. tuberculosis*, are regulated post-translationally by direct interaction of the toxin with its cognate antitoxin protein, which inhibits toxin activity (12, 14, 15, 17). In most cases the genes encoding the antitoxin and toxin are present in a two-gene operon that is autoregulated by antitoxin binding to promoter sequences to repress expression (12). The largest family of Type II TA systems in *M. tuberculosis* is the <u>v</u>irulence-<u>a</u>ssociated <u>p</u>rotein (Vap)BC group, with 50 members (18), the toxins of which are endoribonucleases. Disruption of the antitoxin-toxin interaction results in free toxin in the cell, leading to RNA cleavage and in many, but not all cases, growth arrest. The RNA targets of a number of *M. tuberculosis* VapCs have been identified, with the striking finding that several of these toxins specifically target one or a few tRNAs, thereby precisely altering the *M. tuberculosis* proteome (13, 17, 22–24). Downstream analysis has shown that VapC activity often results in limited and specific effects on the *M. tuberculosis* transcriptome, in addition to effects on the proteome (13, 25, 26).

Though stringent response-mediated antitoxin degradation by Lon or Clp has been suggested to lead to toxin activation in other bacteria (15, 27), for most *M. tuberculosis* TA systems the signals that lead to toxin activation and the mechanism by which these signals are transduced to increase toxin activity are not known. In prior work we identified *in vivo* Ser and/or Thr phosphorylation at 18 unique sites in 11 VapB proteins, including sites in eight VapB antitoxin proteins that showed significantly decreased (>1.9-fold, adj. P<0.01) phosphorylation in a PknA depletion strain (28). We also identified a phosphorylation site in one VapC protein. These findings suggest that reversible Ser/Thr phosphorylation of VapB proteins by receptor Ser/Thr protein kinases may be a novel mechanism for sensing and transducing environmental signals to regulate toxin activity in *M. tuberculosis*. Here, using phosphomimetic and phosphoablative amino acid substitutions in VapB proteins, we provide data for two *M. tuberculosis* VapBC TA systems that show effects of VapB phosphorylation on protein-protein and protein-DNA interactions, and how these substitutions may affect VapC toxin activity.

## Results

### VapB-VapC interaction is decreased when VapB contains phosphomimetic substitutions

Type IIA toxin-antitoxin systems in which the antitoxin VapB directly interacts with its toxin partner VapC and inhibits VapC’s RNA cleavage activity, have been identified in most bacteria, including *M. tuberculosis* (18, 29). In most cases the conditions and signals that modulate the VapB-VapC interaction to allow increased toxin activity are unknown. In a previous phosphoproteomic study, we found that several VapB proteins were Ser/Thr phosphorylated in *M. tuberculosis* H37Rv (28). Moreover, for eight of these, VapB phosphorylation was markedly decreased under PknA depletion conditions. Based on these findings, we hypothesized that Ser/Thr phosphorylation of VapBs may directly impact the interaction between these antitoxins and their cognate VapC toxins.

We selected the VapBC27 and VapBC46 TA systems, both of which showed significantly decreased phosphorylation in the setting of PknA depletion, to study the effect of phosphorylation on VapB molecular interactions and functions. We mutated the phosphoacceptor sites of VapB27 (Thr43) and VapB46 (Ser64) to phosphoablative (Ala) or phophomimetic (Glu for Thr or Asp for Ser) residues. We expressed C-terminus FLAG-tagged VapCs and C-terminus HA-tagged VapBs in *M. smegmatis* and observed inducible expression of both wild type and mutant proteins in the whole cell lysates (Figure 1A & 1B, left panels). To evaluate VapB-VapC interaction, we utilized co-immunoprecipitation (co-IP) with anti-FLAG beads. Western blot analysis of the FLAG-IP revealed substantial binding between wild type and phosphoablative VapB-HAs with their cognate VapC-FLAG (Figure 1A & 1B, right panel).

**Figure 1.**
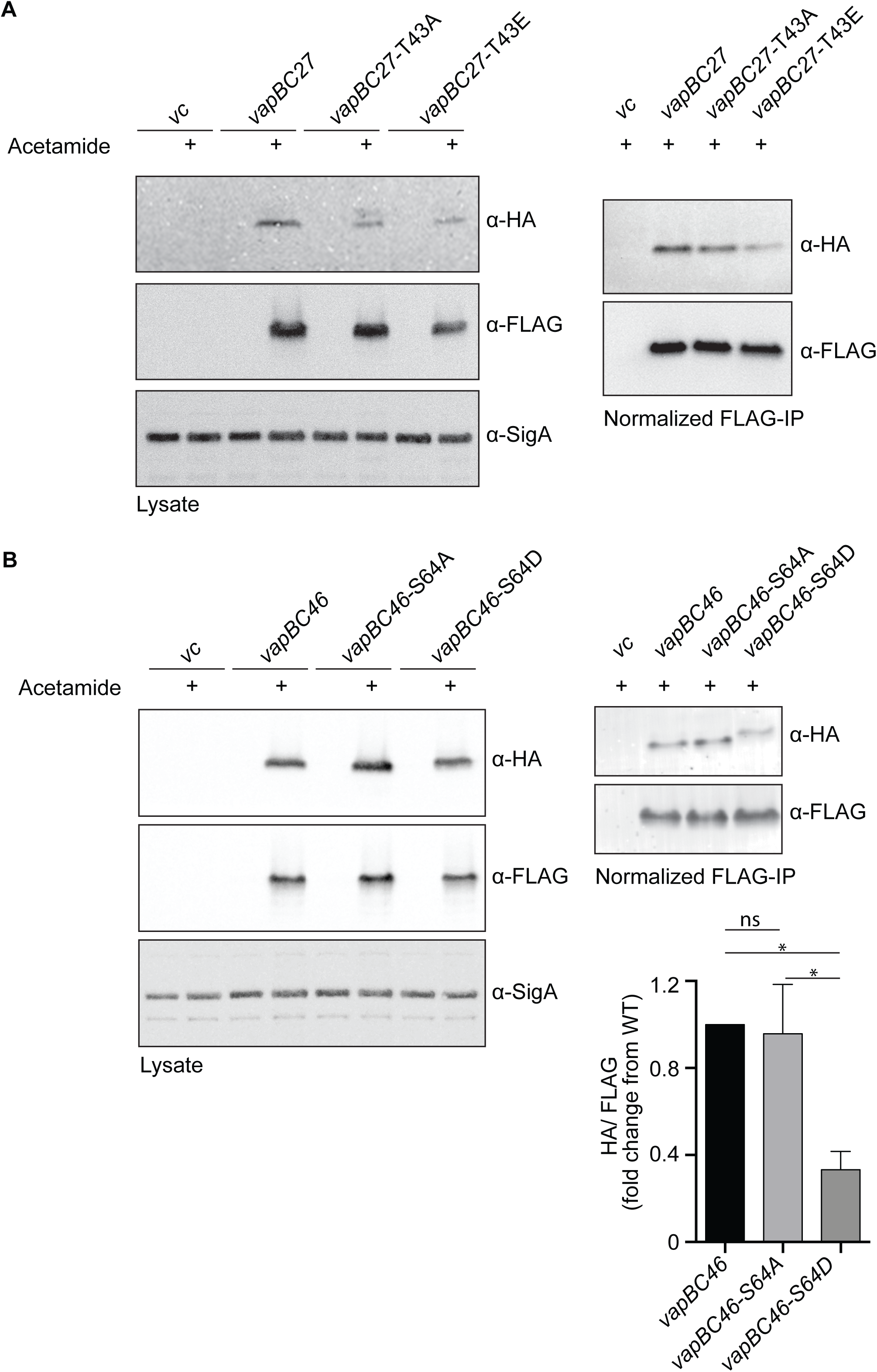
VapB-VapC interaction is negatively regulated by VapB containing phosphomimetic substitutions. **A.** Western blot analysis of *M. smegmatis* mc^2^-155 was performed using whole cell lysates from strains with or without acetamide-induced expression of HA-tagged *M. tuberculosis* VapB27 and FLAG-tagged *M. tuberculosis* VapC27 (left panel). Immunoprecipitation (IP) using an anti-FLAG antibody was performed, and FLAG-VapC-normalized immunoprecipitates were analyzed by western blotting using α-HA and α-FLAG antibodies (right panel). **B.** Western blot analysis of *M. smegmatis* whole cell lysates expressing HA-tagged VapB46 and FLAG-tagged VapC46 was performed as described above (left panel). FLAG-VapC-normalized immunoprecipitates were analyzed by western blotting using α-HA and α-FLAG antibodies (right panel, image). Band intensities were quantified using Li-Cor Image Studio (right panel, bar graph). Asterisk indicates a significant difference (P<0.05 by one way ANOVA) using data from 2 independent experiments. Error bars represent +/- 1 SD.

However, there was a notable decrease in binding between VapB and VapC when the phosphorylation site residue in VapB was mutated to a phosphomimetic residue (Figure 1A & 1B, right panel). These findings suggest that In the context of co-expression of cognate VapB-VapC pairs in the mycobacterial cell, VapBs and VapCs directly interact with each other *in vivo* and that phosphorylation of VapB27 and VapB46 negatively impacts their interaction with their respective VapCs.

### Bacterial two-hybrid assays indicate reduced interaction between *M. tuberculosis* VapC and phosphomimetic-substituted versus native VapB

To further investigate the finding that phosphomimetic mutants of VapB27 and VapB46 show decreased interaction with their cognate VapCs, we conducted Bacterial Adenylate Cyclase Two-Hybrid (BACTH) assays (30). For these experiments, we fused the T18 domain of adenylate cyclase (CyaA) to the C-terminus of wild type and mutant VapBs, and the T25 domain of CyaA to the N-terminus of wild type VapCs. Subsequently, we measured the beta-galactosidase activity in *E. coli* reporter strain BTH101 cells that were co-transformed with a series of VapB-VapC combinations in which the VapB phosphorylation site was modified. Both endpoint and kinetic assays were performed. Our results demonstrated an interaction between wild type and mutant VapBs with wild type VapCs (Figure 2). Consistent with our IP results, in both endpoint and kineteic assays there was no difference between the interactions of the native VapB27 (Figure 2, A-C) and VapB46 (Figure 2, D-E) proteins with their cognate VapC proteins compared to the interactions of the phosphoablative VapB variants. In contrast, we observed a significant decrease in the interaction with the VapB27 and VapB46 phosphomimetic mutants with their VapC partners, again consistent with the findings obtained in the co-IP experiments.

**Figure 2.**
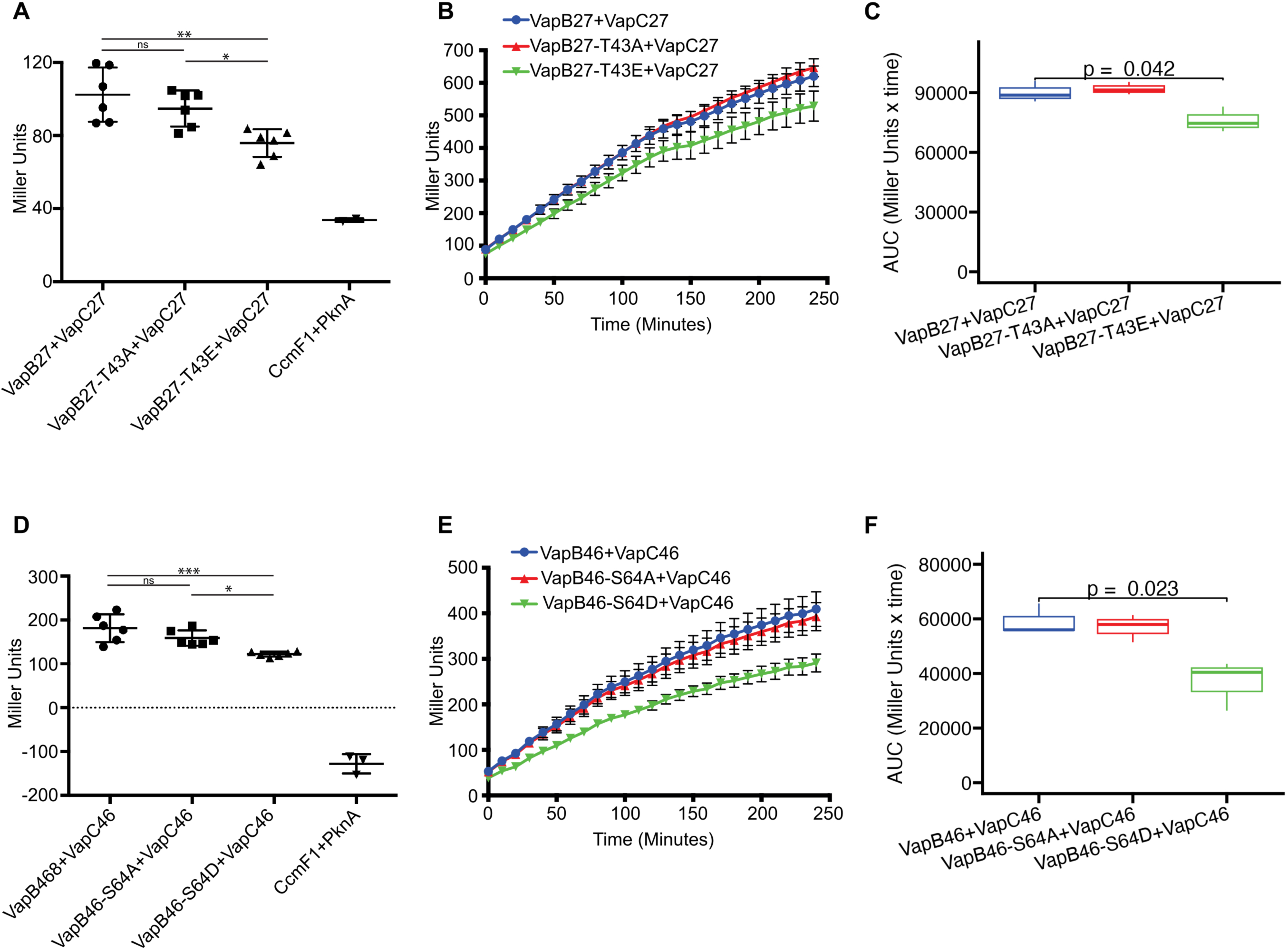
Bacterial two-hybrid assays indicate decreased interaction between *M. tuberculosis* VapC and phosphomimetic-substituted VapB versus native VapB *E. coli* BTH101 cells were co-transformed with plasmids encoding VapB-T18 and T25-VapC fusion proteins, where T18 and T25 are fragments of adenylate cyclase that can form an active enzyme when they interact. The cells were grown on selective media, and the expression of the reporter gene was assayed by β-galactosidase activity. **A-C.** β-galactosidase activity of VapB27-VapC27 in endpoint and time course experiments. **D-F.** β-galactosidase reporter activity of VapB46-VapC46 interaction in endpoint and time course experiments. Data shown for endpoint analysis are representative of two experiments, each with 6 replicates. Statistical analysis of the endpoint assays (Panels **A, D)** was performed by ordinary one-way ANOVA using Graphpad Prism 9. ns, non-significant; *P<0.1; **P<0.01; ***P<0.001. Error bars represent +/ 1 SD. Data shown for the kinetic assays (Panels **B, E**) are representative of two experiments with triplicate cultures. Statistical analysis of kinetic data (Panels **C, F**) was performed using GrowthcurveR (56) and Student’s T-test as described in the Materials and Methods. The T18-CcmF1 and T25-PknA combination was used as the negative control.

### Phosphomimetic substitution in VapB negatively regulates its binding to promoter DNA

It has been shown for a number of TA modules that the *vapB-vapC* operon is subject to autoregulation via binding of the VapB-VapC complex to the promoter region (31). In most cases, this protein-DNA interaction represses transcription of the operon. These data led us to investigate whether the phosphomimetic substitution of VapB also impacts the ability of the VapB-VapC complex to bind to cognate promoter DNA. To this end, we conducted Electrophoretic Mobility Shift Assays (EMSAs) using native VapC, and native or substituted VapB proteins and their corresponding promoter DNA.

We observed clear changes in labelled promoter DNA migration in experiments incorporating the native VapB27 or VapB46 proteins, indicating that the wild type VapB-VapC complexes do indeed bind to their corresponding promoter region in this assay (Figure 3A & 3C). To confirm the specificity of this binding, we performed the reaction in the presence of an excess of unlabeled specific competitor, which led to a reduction in the probe-protein complex intensity and a gradual increase in free probe as the ratio of unlabeled to labeled probe increased (Figure 3B & 3D). Further, we observed a decrease in the probe-protein complex intensity in the VapB27-T43E-VapC27 protein complex reaction (Figure 3A & 3B). For the VapB46-S64D and VapC46 proteins, there was no probe-protein complex formation visible (Figure 3C & 3D).

**Figure 3.**
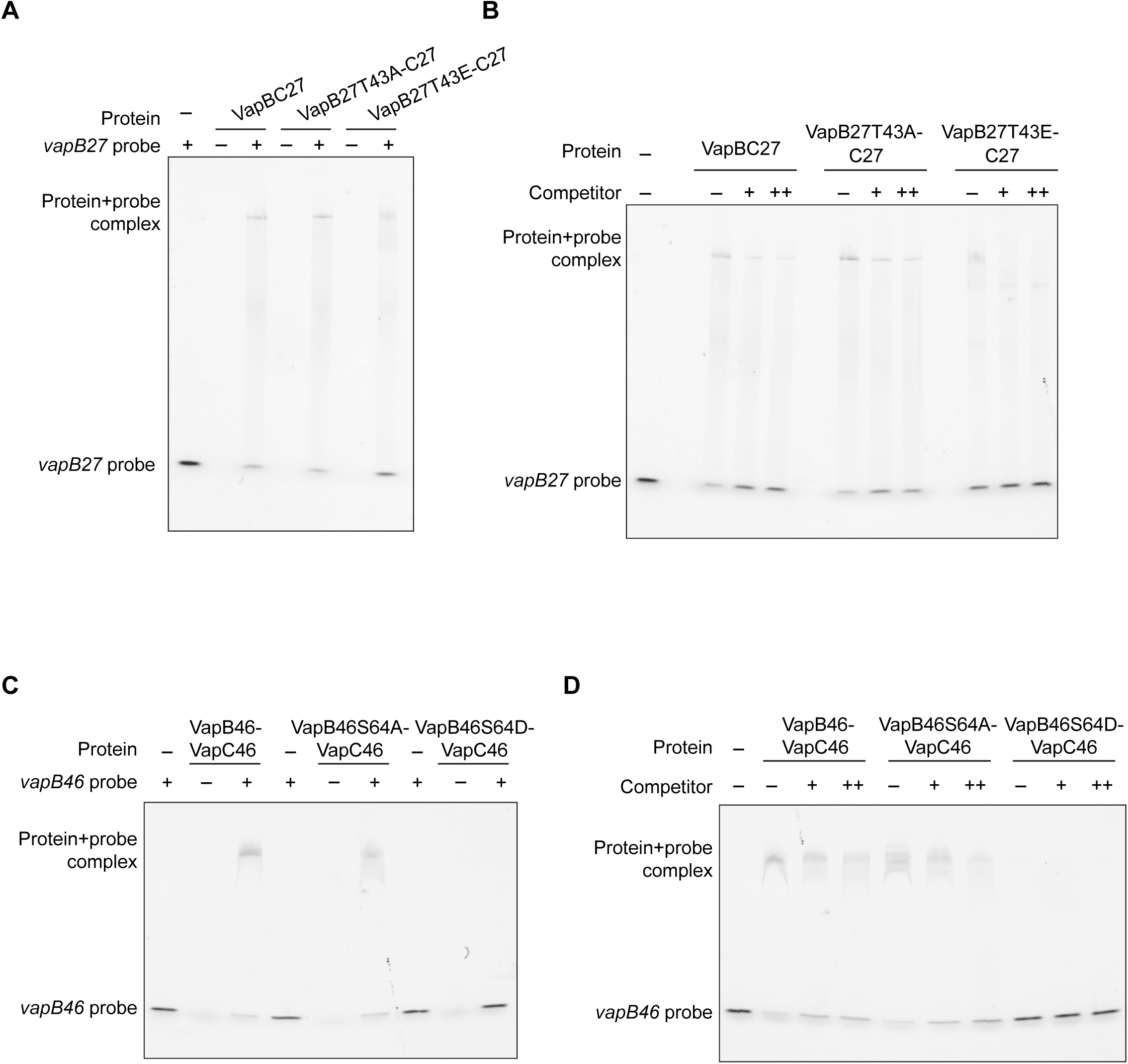
A phosphomimetic substitution in VapB decreases binding of the VapB-VapC complex to promoter DNA. **A.** Electrophoretic Mobility Shift Assays (EMSA) were performed using 200 picomoles of purified VapB27-VapC27, VapB27(T43A)-VapC27, or VapB27(T43E)-VapC27 proteins and 5’-Cy3 labelled *vapB27* promoter DNA as a fluorescent probe. **B.** The assays were performed in the presence of unlabeled specific competitor DNA at probe/competitor molar ratios of 1:25 and 1:50 (+ and ++, respectively). **C.** EMSA was performed using 100 picomoles of purified VapB46, VapB46(S64A), or VapB46(S64D) proteins in combination with 100 picomoles of VapC46, and a 5’-Cy3 labelled *vapB46* promoter DNA as a fluorescent probe. **D.** The assays were performed in the presence of unlabeled specific competitor DNA as described above. Images are representative of 2 to 3 independent experiments.

These findings suggest that VapB phosphorylation interferes with the interaction between the VapB-VapC complex and the *vapBC* operon promoter DNA. This finding may result from direct effects on VapBC complex-DNA binding or from the partial inhibition of formation of the VapB-VapC complex described above, thus affecting its ability to bind to promoter DNA.

### VapC27 is not toxic when expressed in *M. smegmatis* or *M. tuberculosis*

Previous studies have indicated that overexpression of VapC27 does not result in toxicity in *M. smegmatis* (31). Here, we aimed to investigate the effects of phosphoablative or phosphomimetic VapB27 expressed with native VapC27 on growth in *M. smegmatis* and *M. tuberculosis*. To accomplish this, we cloned *vapC27*, *vapBC27* wild type, and *vapBC27* expressing *vapB* phosphorylation site mutants into the anhydrotetracycline-inducible expression vector pMC1s and electroporated the resulting constructs into *M. smegmatis* mc^2^155. We then performed a growth analysis experiment over a period of 24 hours. Consistent with prior reports indicating lack of VapC27 toxicity, we did not observe any significant differences in growth between uninduced and induced cultures across the strains (Figure 4A).

**Figure 4.**
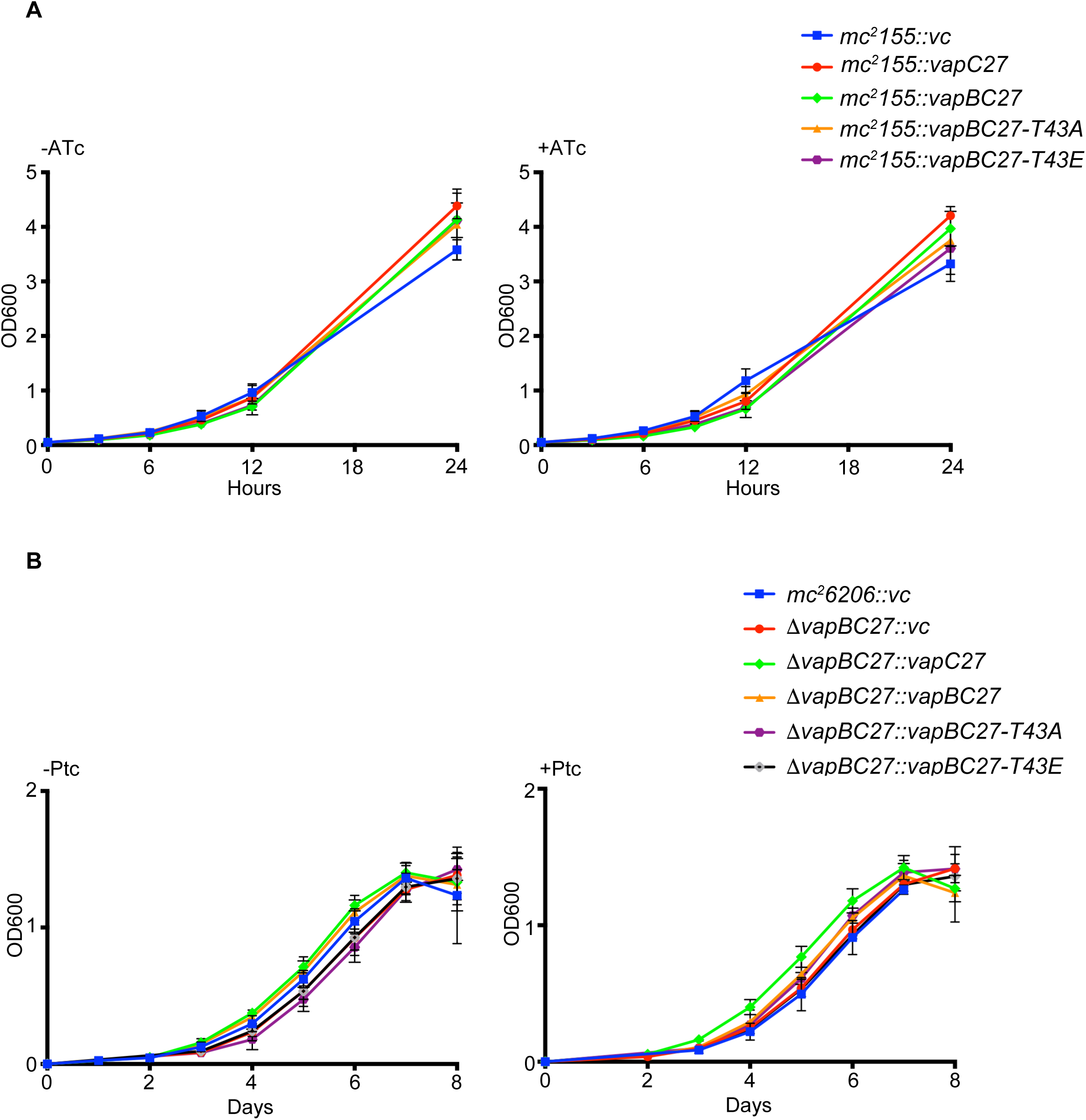
**VapC27 is not toxic when expressed in *M. smegmatis* or *M. tuberculosis*** **A.** *M. smegmatis mc^2^-155* was electroporated with pMC1s-*vapBC27* expressing wild type VapB or with VapB phosphorylation site mutant constructs. Strains were cultured in supplemented Middlebrook 7H9 medium with or without 100 ng/ml anhydrocycline (ATc). Growth was monitored by measuring the OD600 at regular intervals. The data shown are from one experiment with three cultures per strain. **B.** *M. tuberculosis mc^2-^6206ΔvapBC27* was electroporated with pRH2046-*vapBC27* wild type or phosphorylation site mutant constructs and grown in supplemented Middlebrook 7H9 medium with or without 0.5 μg /ml pristinamycin (ptc) inducer. OD600 was recorded daily. The data shown are from one experiment with with three cultures per strain.

To investigate VapC27 toxicity and the effect of VapB variants in *M. tuberculosis* we used the mc^2^6206 auxotroph strain in which we deleted the *vapBC27* operon. This deletion mutant strain was transformed with pristinamycin (Ptc)-inducible *pptr-vapC27* or *pptr-vapBC27* constructs expressing wild type or mutant VapB phosphorylation sites. Again, we did not observe any growth arrest upon overexpression of VapC27 even in the absence of native VapB27, and there were no significant differences in growth between the strains expressing wild type and mutant constructs (Figure 4B). We conclude that VapC27 does not exhibit toxicity when expressed alone or with its cognate VapB antitoxins, under the conditions we tested.

### VapC46 is toxic in *M. tuberculosis* in a manner that varies with co-expression of different *vapB* phosphorylation site alleles

To investigate the impact of *M. tuberculosis* VapC46 on bacterial growth when expressed with native and variant VapB46 proteins, we performed a series of growth experiments. We first electroporated *M. tuberculosis* H37Rv with a Ptc-inducible *pptr-vapC46* construct. We then measured the optical density every 24 hours as well as spotted serial dilutions of uninduced and induced cultures on 7H11 plates. Our results revealed notable differences in the growth dynamics between the induced *vapC46* expression strain and vector control strain, starting from the lag phase of growth (Figure 5A). The observed growth impairment associated with *vapC46* overexpression suggests that VapC46 targets processes that are necessary for the growth of *M. tuberculosis* during infection.

**Figure 5.**
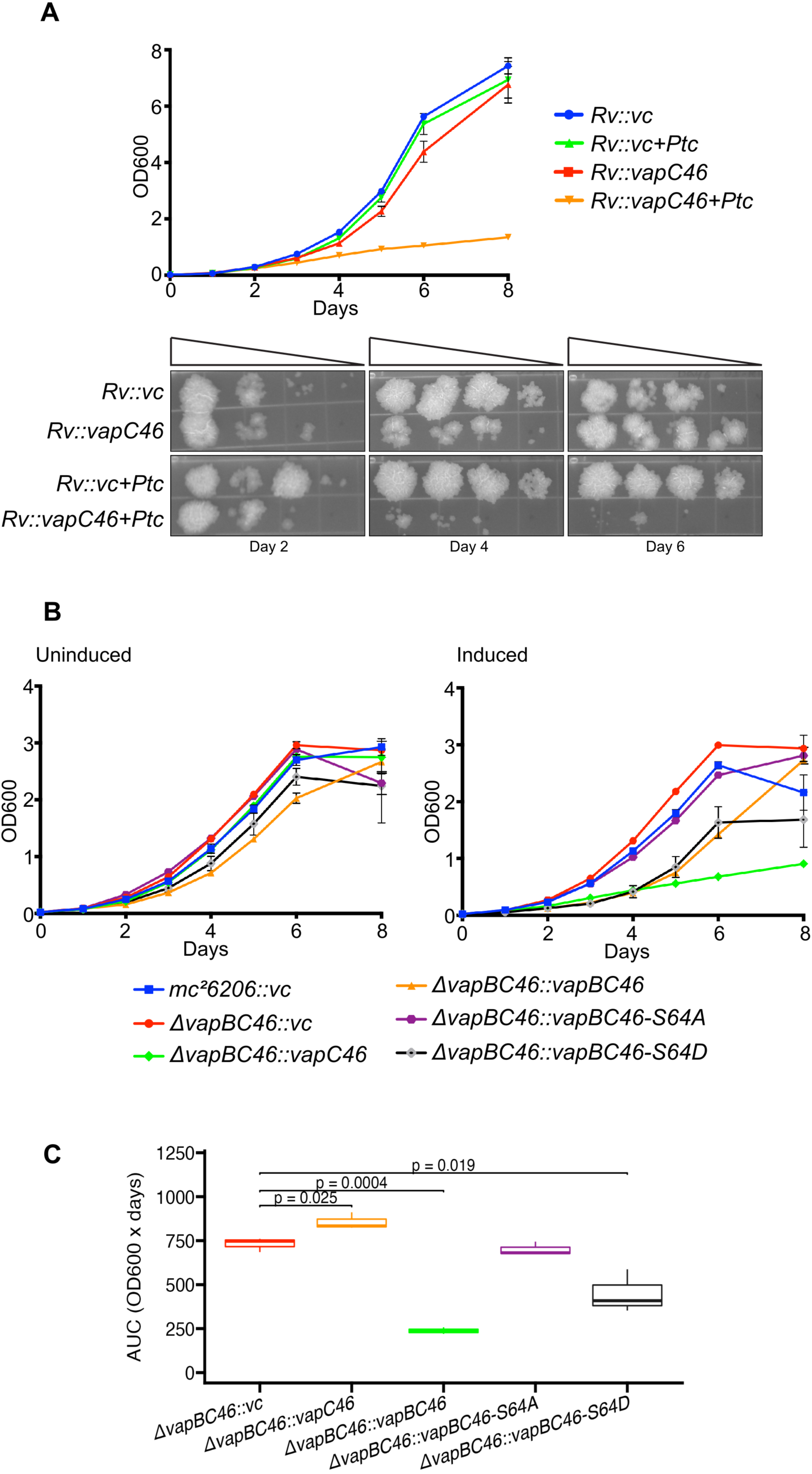
**VapC46 is toxic in *M. tuberculosis* in a manner that varies with co-expression of different *vapB* phosphorylation site alleles** A. *M. tuberculosis H37Rv was* electroporated with the Ptc-inducible vector pRH2046-*vapC46* and grown in supplemented Middlebrook 7H9 medium in the presence or absence of 1 μg/ml Ptc inducer. OD600 was recorded daily. Serial dilutions (10^-1^-10^-4^) of all strains were spotted on 7H9 agar plates on days 2, 4, and 6. B. *M. tuberculosis mc^2-^6206ΔvapBC46 was* co-transformed with pRH2046-*vapC46* and pMind-*vapB46* wild type and phosphorylation site mutant constructs and grown in supplemented Middlebrook 7H9 medium with or without 0.25 μg /ml Ptc and 50 ng/ml tetracycline (tet) inducer. OD600 was recorded daily. The data shown are from one experiment with three cultures per strain. **C.** Box plot comparing AUCs of the growth curve of each strain to the the control *mc^2-^6206ΔvapBC46::vc* (vector control) strain. The central box represents the interquartile range (IQR), spanning from the 25th percentile (Q1) to the 75th percentile (Q3). The thick horizontal line inside the box represents the median AUC value. The whiskers extend from the box to the minimum and maximum non-outlier data points within 1.5 times the IQR. Decreased AUC indicates decreased growth and is a measuare of VapC toxicity.

To investigate the impact on bacterial growth of VapB phosphorylation when co-expressed with native VapC46, we generated a *vapB46-vapC46* deletion mutant in the attenuated *M. tuberculosis* strain mc^2^6206 and introduced inducible *vapC* and inducible alleles expressing wild-type, phosphoablative, or phosphomimetic forms of *vapB46..* As seen in *M. tuberculosis* H37Rv, we observed a significant growth defect in the mc^2^6206 strain expressing *vapC46* alone (Figure 5B, C).

In mc^2^6206 strains containing both native *vapC46* and a variants of *vapB46*, we then induced both *vapB* and *vapC* to to achieve moderately increased expression (1-4-fold relative to mc^2^6206 containing an empty vector) (Figure S1).We observed similar growth of wild type and *vapBC* deletion strains containing the vector control and the strain complemented with *vapB(S64A)-vapC*. In contrast, the deletion strains complemented with wild type *vapB-vapC* and with *vapB(S64D)-vapC* showed substantial growth inhibition during logarithmic growth. Despite this decreased growth over the first several days, the deletion strain complemented with wild type *vapB-vapC* achieved an OD_600_ comparable to wild type at day 8. In contrast, the strain expressing *vapB(S64D)-vapC* did not grow beyond day 6 and had a final OD_600_ that was lower than all other strains except the strain expressing *vapC* alone, which showed minimal growth throughout the time course (Figure 5B, C). These data showing that the *vapB(S64D)-vapC* grows less well than wild type or the deletion mutants complemented with wild type *vapB-vapC* or *vapB(S64A)-vapC* suggest that the decreased binding of VapB(S64D) to VapC that we observed in the co-IP and BACTH assays, and/or the decreased binding of VapB(S64D) to its promoter region, likely occur within the mycobacterial cell. The resulting decreased repression of *vapB(S64D)*-*vapC* and decreased VapB-VapC binding would result in more free VapC, manifested as decreased growth, as seen in these experiments.

### Phosphorylation site substitutions do not inhibit VapB dimerization

Previous studies have demonstrated that VapBs undergo dimerization in solution, and the VapB46-VapC46 complex exists as a heterotetramer or less commonly as a heterooctamer (32). Additionally, VapB dimers have the capability to bind to promoter DNA in the absence of VapC.

To address whether VapB phosphorylation might impair VapB dimer formation, we performed Mycobacterial Protein Fragment Complementation (MPFC) experiments in *M. smegmatis* (33). The results revealed that neither the phosphoablative nor the phosphomimetic phosphorylation site substitutions affected VapB27 or VapB46 dimerization (Figure S2). These findings suggest that impaired VapB dimer formation is unlikely to be the cause for the failure of the phosphomimetic-substituted VapB proteins to bind to the promoter DNA.

## Discussion

The *M. tuberculosis* genome encodes 50 VapBC TA systems, however only a minority of these have been extensively studied. Work to date has shown that several of these TA systems play critical roles in *M. tuberculosis* persistence and stress tolerance, primarily by targeting RNA molecules including mRNA, rRNA and tRNA (22, 24, 34). It has also been shown that under specific stress conditions such as oxidative stress, nutrient deprivation, hypoxia, and nitrosative stress, the expression of specific *vapC* toxin genes is significantly upregulated (17, 25, 35, 36).

Although the increased expression of several *vapCs* in response to stress is well-established the specific mechanisms governing VapC activity, such as their impact on toxin-antitoxin binding and the binding of VapBC to DNA, remain largely unknown. Recent research has suggested a potential regulatory mechanism involving the degradation of VapB antitoxins by proteases such as Clp, Lon or FtsH during stress (37–39). In this model, antitoxin degradation leads to the release of the cognate VapC toxin, which in turn can influence bacterial responses to stress.

In a recent study, we conducted a phosphoproteomic analysis of *M. tuberculosis* in a PknA-depletion strain. Notably, we observed reduced phosphorylation of several VapB antitoxins after depletion of this essential Ser/Thr protein kinase (28). Protein phosphorylation is known to regulate many essential cellular processes, including cell wall synthesis, cell division, metabolism, and virulence (40–46). Therefore, we hypothesized that phosphorylation events might also influence the activity of toxin-antitoxin systems, including VapBCs. To investigate this possibility, we focused on two VapBs, namely VapB27 and VapB46, both of which were phosphorylated *in vivo* and showed decreased phosphorylation in the setting of PknA depletion.

As described in the Results, our initial investigations involved conducting Co-IP experiments using phosphorylation site-substituted VapB proteins. We observed that introduction of the phosphomimetic residue Asp in place of the phosphoacceptor Ser of VapB46 resulted in a major reduction in the interaction between VapB46 and VapC46. In replicate experiments with VapB27-VapC27, we obtained inconsistent results, with evidence of a modest decrease in binding of the phosphomimetic-substituted VapB27 in some but not all experiments.

A distinct approach to investigate VapB-VapC interactions, the *E. coli* BACTH two hybrid assay(30), was consistent with the co-IP results, showing similar interaction of native or phosphoablative VapB27 or VapB46 with their cognate VapCs, but decreased interaction with the phosphomimetic VapBs. As in the co-IP assay, the decrease in VapC binding of the phosphomimetic VapB was greater for the VapB46 interaction compared to VapB27. In both cases the BACTH reporter assays did show substantial residual binding of the phosphomimetic VapB. These findings identify a likely role for VapB phosphorylation in interfering with the VapB-VapC interaction *in vivo* that would result in increased VapC activity, although the mechanism by which the VapB-VapC interaction may be altered by VapB phosphorylation is not known.

The N-terminus of many VapB antitoxins is known to contain a DNA binding domain (47). Analysis of protein family domains using Interpro revealed that the phosphoacceptor site in both VapB27 (T43) and VapB46 (S64) is located near the carboxy-terminal end of their respective DNA binding domains. This led us to investigate the impact of phosphorylation site mutations on promoter DNA binding. In our EMSA experiments, we observed that both wild-type and phosphoablative VapBs, in combination with VapC, exhibited specific binding to the cognate promoter DNA. However, phosphomimetic VapBs showed a clear decrease in binding relative to native or phosphoablative VapBs. To the extent that the VapBC complex acts as a repressor of the auto-regulated vapBC operon, this finding would result in increased *vapBC* transcription and likely increased free VapC protein. In the context of our protein-protein interaction data and published in depth analysis of VapB46-DNA binding (32) this observation suggests that phosphorylation likely regulates both the formation of VapB-VapC oligomers and the binding of this complex to promoter DNA.

Our results showing no effect of VapB phosphorylation on VapB dimer formation in the MPFC assay indicate that phosphorylation-mediated inhibition of VapB dimerization does not explain the diminished DNA binding observed in EMSA experiments that included VapC and VapB variants. Rather, the two factors that we identified may contribute to phosphorylation effects on DNA binding by VapB phosphorylation site variants: the impact of VapB phosphorylation on the VapB-VapC interaction and the decreased ability of the phosphomimetic-substituted VapB proteins that are in a complex with VapC to bind to DNA. Further investigation including exploring other VapB-VapC pairs, are required to comprehensively understand these mechanisms.

Our *in vitro* findings suggest that VapB phosphorylation could have similar effects *in vivo*, with the result that these changes could contribute to increased free VapC abundance and activity under stress conditions *in vivo*. While the absence of VapC27 toxicity in *M. smegmatis* and *M. tuberculosis* precluded our testing this hypothesis for the *vapB-vapC27* TA module, our growth results for strains expressing alleles for native and phosphorylation site mutants of *vapB46* are consistent with the prediction that phosphorylation of VapB46 does lead to increased VapC toxicity in the *M. tuberculosis* cell.

A previous paper described a different post-translational modification (PTM) of a *Salmonella* antitoxin, reversible acetylation, and suggested that this modification regulates cognate toxin activity(48). In that case, the acetylation activity was present in the toxin protein of this Type II TA system. Similar findings for other post-translational modifications of antitoxin proteins in several bacteria, including phosphorylation, adenylation and other PTMs, have been identified or predicted, where the enzyme mediating the PTM is the toxin component of a Type II TA system(49).

In the case of the VapBC systems of *M. tuberculosis*, there is no experimental or bioinformatic evidence that the VapC proteins have intrinsic phosphorylation activity. Rather, there are 11 Ser/Thr protein kinases (STPK) in this organism, of which 9 are single-pass transmembrane proteins. This structure is typical of receptor-type protein kinases that sense and transduce extracytoplasmic environmental signals by phosphorylation of cytoplasmic proteins. This observation suggests that the *M. tuberculosis* STPKs play a major role in phosphorylation of VapB proteins in *M. tuberculosis*, providing a mechanism by which toxin activity can be regulated by extracytoplasmic stresses. This interpretation is supported by the markedly decreased phosphorylation of several VapB proteins that we previously identified when PknA was depleted in *M. tuberculosis* cells (28). The recent identification of differential phosphorylation of over half of all *M. tuberculosis* VapB proteins, as well as several VapCs, in response to overexpression or deletion of one or more STPKs in *M. tuberculosis* further highlights the importance of this mechanism for regulating *M. tuberculosis* physiology (50). At this time, however, for most *M. tuberculosis* VapC toxins the extracytoplasmic and intracellular environmental signals and mechanisms of signal transduction that lead to increased toxin expression and toxicity are unknown.

In summary, our results identify two possible mechanisms by which VapB phosphorylation can affect VapC toxin activity. First, using phosphomimetic and phosphoablative substitutions in two VapB proteins, we have shown that phosphorylation interferes with VapB binding to VapC. Second, we have shown that VapB phosphorylation interferes with its binding to the promoter region of the cognate *vapBC* operon. Both of these effects are predicted to increase free VapC in the cell, and thus increase VapC toxicity. This prediction is supported by our observation of slower growth of *M. smegmatis* and *M. tuberculosis* strains containing a *vapBC46* construct that expresses phosphomimetic VapB, compared to growth of strains containing *vapBC46* constructs that express native or phosphoablative VapB. These data support the concept that toxin activity in *M. tuberculosis* can be rapidly altered by antitoxin phosphorylation/dephosphorylation-mediated signal transduction to allow adaptation of this pathogen to changing environments during infection.

## Materials and Methods

### Strains, media, and reagents

*M. tuberculosis* H37Rv (ATCC 27294), attenuated *M. tuberculosis* mc^2^ 6206 (H37RvΔ*leuCD*Δ*panCD*, a kind gift of Dr. William Jacobs Jr laboratory) (51), and *M. smegmatis* mc^2^155 (ATCC 700084) were used as wild type mycobacterial strains. mc^2^6206Δ*vapBC27* was kindly provided by Dr. Nancy Woychik’s lab. *E. coli* Top10 (Invitrogen) was used for all cloning experiments while *E. coli* BL21 (DE3) Codon Plus (Stratagene) cells were used for purification of recombinant proteins. *M. smegmatis* (mc^2^155) and *M. tuberculosis* (H37Rv) strains were grown in Middlebrook 7H9 media (Difco) supplemented with 10% ADN supplement (albumin, glucose, and sodium chloride) at 120 rpm at 37°C with appropriate antibiotics, when needed. The media was supplemented with 50 μg/ml pantothenic acid and 100 μg/ml leucine for the attenuated mc^2^6206 strain. Pristinamycin 1A (Ptc) obtained from Cayman Chemical, Canada. Acetamide, tetracycline (Tet) and anhydrotetracycline (ATc) were purchased from Sigma. Restriction endonucleases and other enzymes were obtained from New England Biolabs.

### Co-Immunoprecipitation

Point mutations of VapB phosphoacceptor residues (VapB27-T43→A and T43→E; VapB46-S64→A and S64→D) were generated using the NEB Q5 mutagenesis kit. PCR amplicons of *vapB* and *vapC*s were cloned into pRH2790 and pRH2017 vectors, respectively. In these constructs, VapBs would have a C-terminal HA tag, and VapCs would have a C-terminal 3X-FLAG tag. pRH2790-*vapB* wild type and mutant constructs in combination with pRH2017-*vapC* were electroporated into *M. smegmatis* mc^2^155 strain. Resultant colonies were inoculated in 7H9 media and grown till OD_600_ reached ∼0.3. They were then induced with 0.2% acetamide and grown for another 6 hrs. Whole cell lysates were prepared as described earlier (52). FLAG-tagged VapCs were immunoprecipitated using anti-FLAG sepharose beads (Cell Signalling Technologies). Immunoprecipitated samples were then processed for western blotting.

### Western blot analysis

For western blotting, protein concentrations were measured using the BCA assay and 20 μg of proteins from whole cell lysates were resolved on pre-cast 12% tris-glycine PAGE (Bio-Rad) followed by transfer to a LF-PVDF membrane using a Bio-Rad semi dry transfer apparatus. The membranes were then probed with appropriate dilutions of α-FLAG (Sigma, 1:4000), α-HA (Invitrogen, 1:1000), or α-Sigma70 (Neogen, 1:2000) antibodies. Incubation with primary antibody was followed by incubation with HRP-linked anti-mouse/rabbit secondary antibody (Cell Signalling Technology). An Azure c600 gel imaging system was used to visualise bands. For imaging with LI-COR, after incubation with primary antibody, membranes were incubated with IRdye 800CW-conjugated secondary antibodies (LI-COR). An Odyssey Fc imager was used to visualise bands and bands were quantified using Image Studio software.

### Bacterial Adenylate Cyclase Two-Hybrid (BACTH) assays

BACTH assays were used to investigate interaction between VapC and wild type and phosphomutant VapBs (30). VapBs and VapC were expressed as C-terminus and N-terminus fusion proteins to the T18 and T25 fragments of *Bordetella pertussis* adenylate cyclase (CyaA) in *E. coli* BTH101 cells, respectively. pUT18-*vapB* and pKT25-*vapC* constructs were co-transformed into *E. coli* BTH101 cells and plated onto LB agar containing appropriate antibiotics (Ampicillin 100 μg/ml and Kanamycin 50 μg/ml), 0.5 mM IPTG and 40 μg/ml X-gal. Resulting blue colonies were inoculated into fresh LB media with antibiotics and 0.5 mM IPTG and were grown with continuous shaking at 200 rpm at 30°C until OD_600_ reached ∼0.5-0.6. The tubes were chilled on ice for 20 min and diluted with PM2 assay media (70 mM Na_2_HPO_4_.12H_2_0, 30 mM NaH_2_PO_4_ H_2_0, 1 mM MgSO_4,_ 0.2 mM MnSO_4,_ pH 7.0, 100 mM β-mercaptoethanol). 60 μl chloroform and 30 μl 0.1% SDS were added per ml of diluted cells and the tubes were vortexed vigorously to permeabilize the bacterial cells. Tubes were equilibrated in a 28°C water bath for 10 min. Aliquots were transferred into a 96-well UV-transparent flat-bottom plate. The enzymatic reaction was started by adding 0.4% o-nitrophenol β-galactoside (ONPG) pre-equilibrated at 28°C. Kinetic assays were performed in a microplate reader set at 28°C measuring OD_420_ every 10 min for 4 hrs. For endpoint assays, the reaction was stopped by adding 1M Na_2_CO_3_ after yellow color developed in the positive control well. OD_420_ was measured in a microplate reader.

### Production of recombinant VapBCs and Electrophoretic Mobility Shift Assay (EMSA)

VapBC27-His, VapB46-His and VapC46-His wild type and mutant proteins were purified from *E. coli* BL21 (DE3) Codon plus cells as previously described (23). Binding of the VapB-VapC complex to the *vapB* promoter region was evaluated by EMSA using a 5’-Cy3 labelled *vapB* promoter oligo (vapB27: 5’ cggtgagtcgacttccgcacctgagtgggaagccattggtatcgtattcccatga 3’; vapB46: 5’ tcctcccagctcagcgccaaccaaccgaggaacaacgcgccgactttttcag 3’). These oligos were annealed to their respective 5’-Cy3-labelled complementary oligo to form a double-stranded probe. The probe:protein molar ratio used for the VapBC27 complex was 1:1000 while it was 1:500 for VapB46 and VapC46. The reaction buffer consisted of 20 mM Tris-HCl (pH 7.5), 150 mM KCl, 2 mM MgCl_2_, 1 mM dithiothreitol (DTT), 5 µM EDTA, 5% glycerol, 0.05% NP-40, 50 ng poly(dI). poly(dC), and 200 fmol labelled probe. The samples were incubated for 30 min at room temperature and then resolved on a 7.5% TGX Stain free gel (Bio-Rad) at 120 V at 4°C for 1 hr. The gel was scanned using an Azure c600 gel imaging system. Unlabelled specific competitor DNA was used at 1:25 and 1:50 molar ratios (labelled:unlabelled), when required.

### Generation of a vapBC46 deletion mutant

The *vapB46-vapC46* operon (Rv3385c-Rv3384c) in *M. tuberculosis* mc^2^6206 strain was deleted using Oligonucleotide-mediated Recombineering followed by BxB1 Integrase Targeting (ORBIT) method (53). The Bxb1 *attP*-containing oligonucleotide was flanked by the first and last 30 nucleotides of the *vapB46-vapC46* operon. To confirm successful deletion, we performed PCRs using oligonucleotides targeting the antitoxin-toxin module and from internal flanking regions in the PKM464 plasmid.

### Growth kinetics experiments

To perform growth experiments in *M. smegmatis*, *vapC27* alone or *vapBC27* expressing native or phosphorylation site VapB substitutions were cloned into the ATc-inducible pMC1s vector (54). They were electroporated into *M. smegmatis* mc^2^155 and resultant colonies were inoculated in 7H9 media supplemented with 10% ADN with 25 μg/ml kanamycin. Strains expressing VapC or VapBCs were induced by adding ATc at a final concentration of 100 ng/ml. The cultures were grown with continuous shaking at 120 rpm at 37°C. Growth was monitored by recording OD_600_ over 24 hrs. For growth experiments in *M. tuberculosis,* constructs expressing *vapC27* alone or *vapBC27* wild type or *vapB* phosphomutants were cloned into Ptc-inducible pRH2046 vector and were electroporated in*. M. tuberculosis* Mc^2^6206Δ*vapBC27* strain.

Resultant colonies were inoculated in 7H9 media with appropriate supplementation and 100 μg/ml spectinomycin. Expression of VapC27 or VapBC27 was induced by adding Ptc at a final concentration of 0.5 μg/ml. The cultures were grown with continuous shaking at 120 rpm at 37°C..

To evaluate VapC46 toxicity, pRH2046-pptr-*vapC46* or the empty vector control was electroporated in *M. tuberculosis* H37Rv (wild type) and resultant colonies were inoculated in supplemented 7H9 medium with spectinomycin. VapC46 was induced by adding 1 μg/ml Ptc and cultures were grown at 120 rpm at 37°C. Growth was monitored over a period of one week by measuring the OD_600_ of the liquid cultures as well as observation of serial dilutions of cultures (10^-1^-10^-4^) spotted on 7H9 agar plates. To assess the effect of *vapB* phosphomutants on VapC toxicity, constructs expressing native VapB46 or VapB46 phosphorylation site substitutions were cloned into Tet-inducible pMind constructs and electroporated into *M. tuberculosis* mc^2^6206Δ*vapBC46* along with pRH2046-pptr-*vapC46* (55). Resultant colonies were grown in

7H9 medium with spectinomycin (100 μg/ml) and zeocin (25 μg/ml). VapC46 and VapB46 were induced by adding 0.25 μg/ml Ptc and 50 ng/ml Tet, respectively. Growth in liquid medium was monitored by recording OD_600_ every 24 hrs for one week. Statistical analysis to compare growth of mc^2^6206Δ*vapBC46* complemented by the empty vector to mc^2^6206Δ*vapBC46* complemented by *vapC* alone or by vapC plus the *vapB* variants was performed using Growthcurver software (56) by comparing their area under the curve value (auc_l) by Student’s t-test.

### Total RNA isolation and quantitative RT-PCR

To determine the expression levels of inducible *vapB* and *vapC* constructs, samples were collected 48 hrs after addition of inducer, resuspended in Trizol and mechanically disrupted with 0.1 mm zirconia beads in a bead beater. Total RNA was extracted using Direct-zol RNA Miniprep kit (Zymo Research) according to the manufacturer’s protocol. RNA was treated once more with Turbo DNase (Thermo Fisher) and the quantity and quality were checked by taking absorbance at 260 and 280 nm using a Nanodrop instrument. Reverse transcription of total RNAs was carried out using Quanta Biosciences qScript cDNA synthesis kit. Absence of contaminating genomic DNA was confirmed by performing qPCR of RNA without reverse transcription (No template control). Quantitative RT-PCR of cDNAs was performed using Maxima SYBR Green/ROX qPCR Master Mix (Thermo Scientific) on a QuantStudio 3 real time PCR system. Data were analyzed using ΔΔCt method with *sigA* as areference gene (57).

## Data availability statement

All data related to the findings in this article are presented in the main text and figures, and in the Supplemental Materials file.

## Acknowledgments

We thank the William Jacobs laboratory (Albert Einstein College of Medicine) for providing the *M. smegmatis* strain mc^2^-155 and the attenuated H37Rv-derived *M. tuberculosis* strain mc^2^-6206 (Δ*panCD* Δ*leuCD*). We thank Brenda Nguyen and Jolie Ren for technical support in protein purification and molecular cloning. Funding for this research was provided by the National Institutes of Health grants R01AI154464 to NW and R21AI146539 to RNH. The funders had no role in study design, data collection and interpretation, or the decision to submit the work for publication.

## Notes

### Competing Interest Statement

The authors have declared no competing interest.

