## Supplementary material for "Phosphorylation of VapB antitoxins affects intermolecular interactions to regulate VapC toxin activity in *Mycobacterium tuberculosis*": Suppl figs 1 & 2 and legends

Figure S1

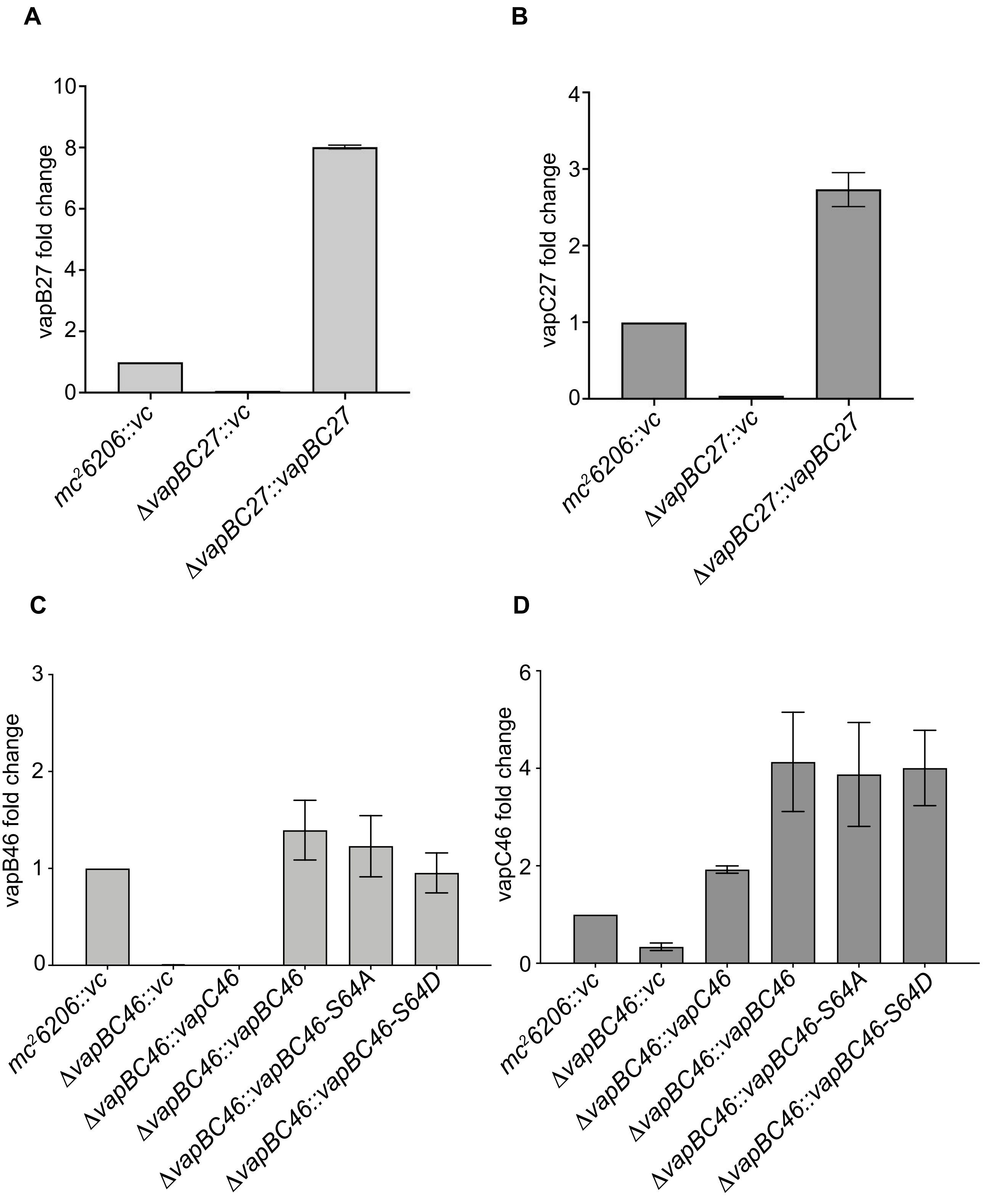
A
vapB27 fold change
0
2
4
6
8
10
mc²6206::vc
ΔvapBC27::vc
ΔvapBC27::vapBC27
B
vapC27 fold change
0
1
2
3
4
mc²6206::vc
ΔvapBC27::vc
ΔvapBC27::vapBC27
C
vapB46 fold change
0
1
2
3
mc²6206::vc
ΔvapBC46::vc
ΔvapBC46::vapC46
ΔvapBC46::vapBC46
ΔvapBC46::vapBC46-S64A
ΔvapBC46::vapBC46-S64D
D
vapC46 fold change
0
2
4
6
mc²6206::vc
ΔvapBC46::vc
ΔvapBC46::vapC46
ΔvapBC46::vapBC46
ΔvapBC46::vapBC46-S64A
ΔvapBC46::vapBC46-S64D

Figure S2

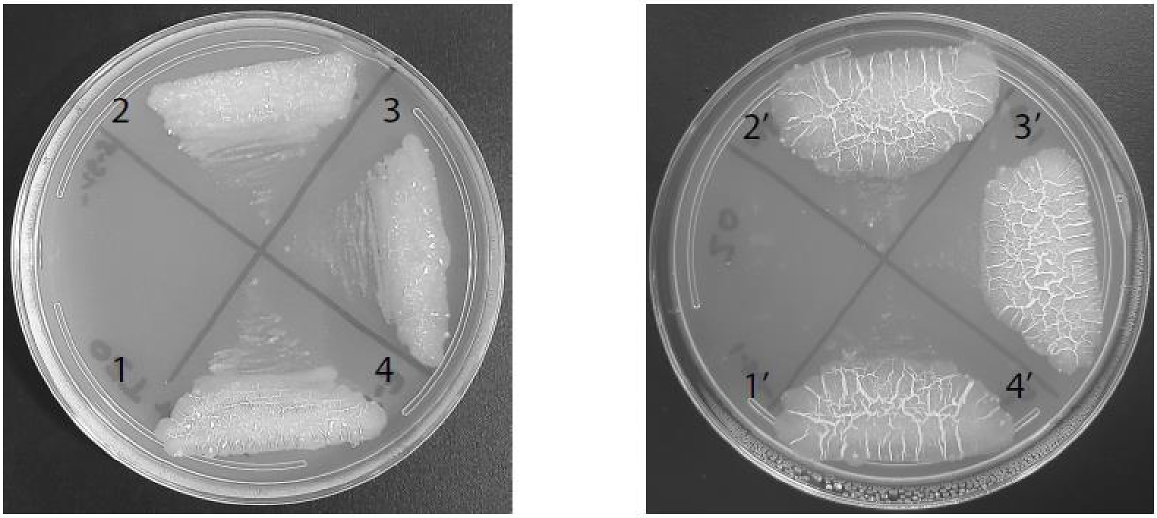


1. VapB27-F[1, 2]+F[3]
2. VapB27-F[1, 2]+VapB27-F[3]
3. VapB27-T43A-F[1, 2]+VapB27-T43A-F[3]
4. VapB27-T43E-F[1, 2]+VapB27-T43E-F[3]

1. VapB46-F[1, 2]+F[3]
2. VapB46-F[1, 2]+VapB46-F[3]
3. VapB46-S64A-F[1, 2]+VapB46-S64A-F[3]
4. VapB46-S64D-F[1, 2]+VapB46-S64D-F[3]

**Figure S1.** Expression of induced VapB and VapC in *M. tuberculosis* $mc^2 6206$ strains.

Total RNA was isolated from all strains 48 hours post induction as described in Materials and Methods. Briefly, *M. tuberculosis* $mc^2 6206\Delta vapBC27$ strains electroporated with wild type or phosphorylation site mutant constructs were induced with 0.5 µg /ml Ptc inducer. In *M. tuberculosis* $mc^2 6206\Delta vapBC46$ strains, vapB46 and vapC46 were induced with 50 ng/ml tet and 0.25 µg /ml Ptc, respectively. Equal amounts of RNA were used to set up quantitative RT-PCR. **A-B.** Expression of vapB27 and vapC27 as determined by qRT-PCR. **C-D.** Expression of vapB46 and vapC46 as determined by qRT-PCR.

**Figure S2**. MPFC assays showing VapB homodimer formation

*M. smegmatis* $mc^2 155$ was electroporated with MPFC-F[1,2] and MPFC-F[3] plasmids expressing wild type and mutant versions of VapB27 or VapB46. Transformant colonies were inoculated in 2 ml Middlebrook 7H9 supplemented media. 10 µl of culture were streaked onto 7H11 agar plates containing 30 µg/ml trimethoprim and incubated at 37°C for 3-4 days. Growth on trimethoprim-containing media indicates that the two proteins interact with each other.
